# Diverse Database and Machine Learning Model to Narrow the Generalization Gap in RNA Structure Prediction

**DOI:** 10.1101/2024.01.24.577093

**Authors:** Albéric A. de Lajarte, Yves J. Martin des Taillades, Colin Kalicki, Federico Fuchs Wightman, Justin Aruda, Dragui Salazar, Matthew F. Allan, Casper L’Esperance-Kerckhoff, Alex Kashi, Fabrice Jossinet, Silvi Rouskin

## Abstract

Understanding macromolecular structures of proteins and nucleic acids is critical for discerning their functions and biological roles. Advanced techniques—crystallography, NMR, and CryoEM—have facilitated the determination of over 180,000 protein structures, all cataloged in the Protein Data Bank (PDB). This comprehensive repository has been pivotal in developing deep learning algorithms for predicting protein structures directly from sequences. In contrast, RNA structure prediction has lagged, and suffers from a scarcity of structural data. Here, we present the secondary structure models of 1098 pri-miRNAs and 1456 human mRNA regions determined through chemical probing. We develop a novel deep learning architecture, inspired from the Evoformer model of Alphafold and traditional architectures for secondary structure prediction. This new model, eFold, was trained on our newly generated database and over 300,000 secondary structures across multiple sources. We benchmark eFold on two new test sets of long and diverse RNA structures and show that our dataset and new architecture contribute to increasing the prediction performance, compared to similar state-of-the-art methods. All together, our results reveal that merely expanding the database size is insufficient for generalization across families, whereas incorporating a greater diversity and complexity of RNAs structures allows for enhanced model performance.

## Introduction

The structural integrity of RNA is often critical to its function, playing a vital role in how RNA molecules interact within the cell. While mRNA’s primary sequence is decoded in function (e.g. translation), RNA harbors a second layer of information—its ability to form secondary structures. These structures can conceal or expose crucial regulatory binding sites for proteins, microRNAs, and other RNAs [1., 2., 3., 4.]. While X-ray crystallography, Nuclear Magnetic Resonance (NMR), and CryoEM are adept at elucidating RNA’s tertiary structure, their applicability is limited for long RNA sequences (>200nt). Moreover, it is plausible that many long RNAs, apart from specialized ones such as ribosomal RNAs, may not adopt stable tertiary structures, further complicating their studies with conventional methods. As a result, the focus shifts to RNA’s secondary structure, which is an intrinsic characteristic of all RNA molecules. To predict this secondary structure, dynamic programming algorithms are frequently used, identifying the conformation with the lowest energy based on the Turner nearest neighbor rules [5., 6.], a set of experimental parameters for the energy of different base pairs in the context of diverse structure motifs.

Although widely used, these algorithms have their limitations, typically confined to canonical base pairs and structures and often unable to predict more complex interactions such as non-canonical base pairs or pseudoknots. A contrasting approach is to learn RNA structure models directly from data. In this vein, recent advancements have seen the emergence of numerous deep learning methods [7., 8., 9.]. This methodology makes minimal assumptions about the thermodynamic rules governing RNA structure and, with adequate data, could capture more complex motif types. Nonetheless, a significant challenge in deep learning is ensuring a sufficiently diverse and high-quality dataset. A training set lacking in diversity results in poor performance in out-of-domain contexts. This problem is exacerbated in RNA studies due to the unique sequences and structures of different RNA families, leading to many distinct domains. For instance, Flamm et al. [10.] demonstrated how performance significantly declines when algorithms are trained on a specific set of families or sequence lengths and then tested on different ones. Similarly, Bugnon et al. [11.] examined this effect, observing diminished performance when applying published algorithms to very long ncRNA and mRNA. Szikszai et al. [12.] also highlighted the risks of using test sets composed of similar families and advocated for the curation of inter-family test sets.

There are a few databases commonly used by all deep learning methods. The bpRNA database [13.] consists of over 100,000 sequences and structures aggregated from multiple sources including PDB and the RNA families database (Rfam). The PDB database [14.] includes 1,790 RNA tertiary structures as of December 2023. ArchiveII [15.] and RNAStralign [16.] contain 3,975 and 30,451 sequences, respectively. Additionally, a substantial dataset of 806,000 sequences, together with chemical probing data for DMS and SHAPE and data-derived structure models, was recently released for the Ribonanza competition [17.]. Nevertheless, many of these datasets require extensive filtering to eliminate redundant and low-quality sequences. For example, bpRNA90 is a curated subset of bpRNA featuring 28,000 non-redundant sequences, which is 27% of the original database. Additionally, these databases tend to be biased towards a limited number of short ncRNA families, which may adversely affect the generalization capabilities of commonly used algorithms.

We examine the generalization ability of four widely-used secondary structure algorithms, each employing different methodologies. RNAStructure Fold [18.] is a classic model that utilizes the Nearest Neighbors parameters and dynamic programming to determine the minimum free energy of a structure. EternaFold [19.], adapted from ContraFold [20.], employs context-free grammar to learn the probability of different motifs and structures. MXFold2 [8.], a deep learning model, predicts the probability of all potential base pairs and conformations as multiple base pairing matrices. Both EternaFold and MXFold2 are hybrid approaches, leveraging dynamic programming to determine the minimum free energy structure from the predicted energy motifs. UFold [7.], on the other hand, predicts a base pairing matrix from the sequence using only a neural network (“end-to-end model”) outfitted with a minor post-processing step for conversion into a valid structure. We benchmark these algorithms on a set of challenging sequences. The results indicate poor generalization capabilities. This finding led us to develop a new database of relevant and diverse sequences in an effort to mitigate the generalization gap.

The state-of-the-art method for high throughput RNA structure prediction involves probing the RNA molecule with reagents that measure the probability of each nucleotide being paired or unpaired. The two most frequently used methods are DMS-MaPseq and selective 2′-hydroxyl acylation analyzed by primer extension (SHAPE). In DMS-MaPseq dimethyl sulfate (DMS) reacts at the Watson-Crick face of adenine and cytosine, while SHAPE identifies the most flexible parts of the RNA backbone, typically where unpaired bases are found. Both methods induce mutations in the sequence during reverse transcription, which can be detected through high-throughput next-generation sequencing. Analyzing a multitude of sequencing reads provides a mutation fraction per nucleotide which, after normalization, is interpreted as a pairing probability. This data, when added as a pseudo-energy term to classical algorithms, significantly enhances the accuracy of the predicted structure [21., 22.]. Here, we employed the DMS-MaPseq method to probe and model the structure of a variety of RNAs from new and underrepresented families. Our efforts aim to improve generalization across different RNA domains. The data collected, covering biologically significant sequences such as the 3’ ends of mRNAs and pri-miRNAs of the human genome, is now accessible in our online database (https://rnandria.org/). Using this new data along with existing RNA structure databases, we trained a new architecture, eFold, based on the Evoformer from Alphafold [23.] and Convolutional Neural Networks. We curate a set of challenging test structures and demonstrate enhanced generalization capabilities compared to state-of-the-art deep learning models.

## Results

### Secondary structure algorithms do not generalize across different types of RNAs

Current secondary structure predictions rely on the same databases for their training sets. A notable issue is the lack of diversity in these datasets, which are predominantly biased towards short ncRNAs. Rfam [24.] is a database of all the known RNA families and clans, counting 146 clans to date, and consistently adding new ones. However, only a few clans such as tRNA, rRNA and sRNA are significantly represented in commonly used databases, and most sequences are typically less than 200 nucleotides long (Figure 1a, Supplementary Figure 1a-d). This bias allows deep learning models to excel within these family domains, but their performance substantially diminishes outside of them [10.,11.,12.].

**Figure 1:**
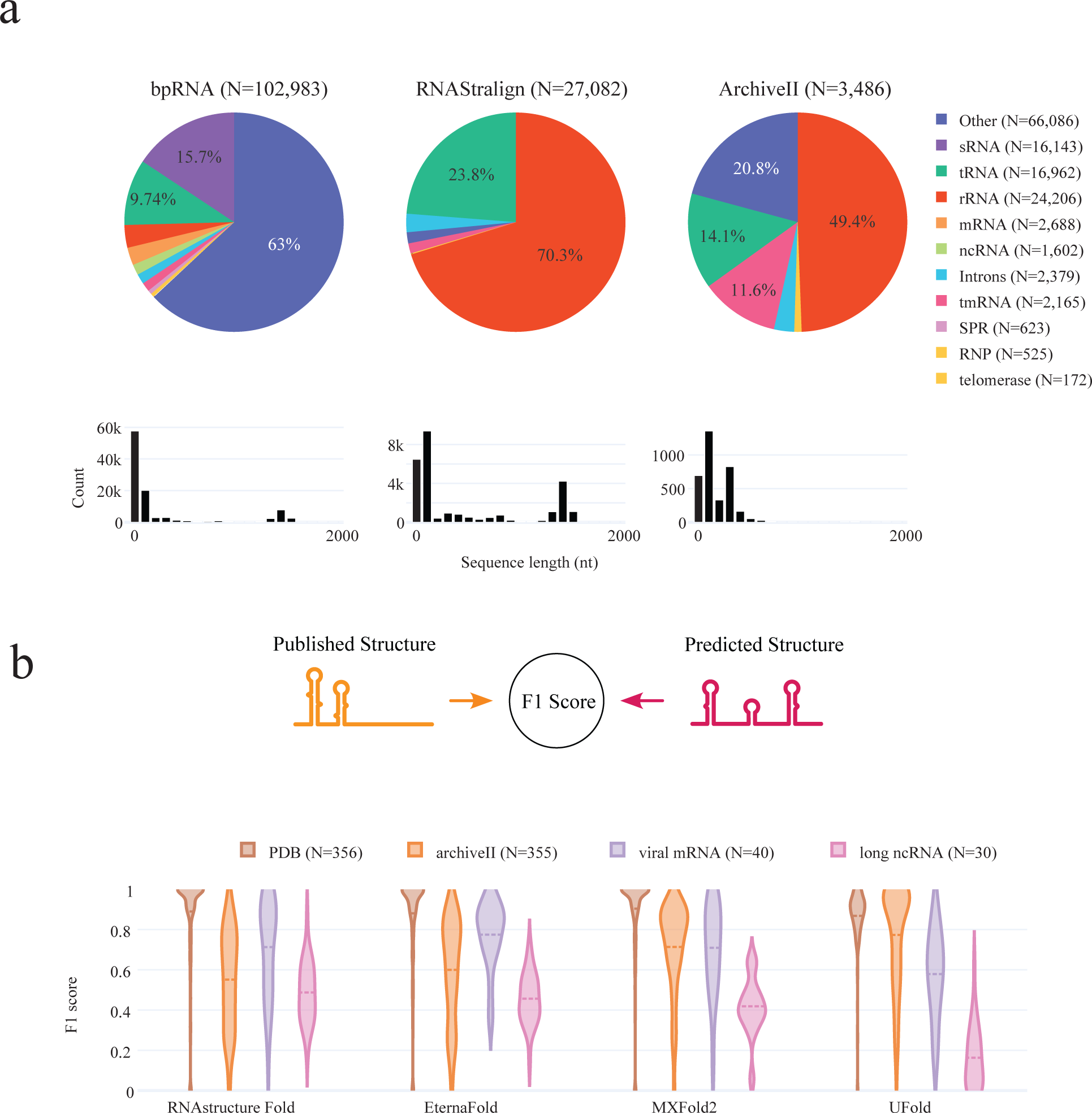
Performance of state-of-the art algorithms on diverse RNA domains. **(a)** RNA types and sequence length distribution of bpRNA, RNAstralign, and ArchiveII datasets. **(b)** A violin plot illustrating the distribution of F1 scores comparing the predicted structure model by each algorithm against a published structure model across four distinct test sets: PDB, ArchiveII, Viral mRNAs, and Long ncRNAs. The F1 score, calculated as the harmonic mean of precision and sensitivity, spans from 0 to 1.

To evaluate performance beyond common testing sets, we curated four test datasets from published RNA structures. The first dataset comprises RNA structures from the Protein Data Bank (PDB), consisting of 356 sequences obtained by excluding entries with multiple chains, such as RNA-RNA, RNA-DNA, or RNA-protein complexes, resulting in a collection of predominantly short noncoding RNAs. Our second dataset, ArchiveII, a commonly used test set in previous studies [7., 8., 9.], was refined by removing all sequences also present in bpRNA and RNAstralign, as these databases are frequently utilized as training sets. The final two test sets aim to evaluate performance across new families and various sequence lengths. The long noncoding RNA dataset (Long ncRNA), curated by Bugnon et al. [11.], includes 30 long RNA sequences ranging from 1000 to 2000 nucleotides, with structures exceeding 2000 nucleotides segmented into smaller modular structures, as detailed in the Methods section. The fourth dataset consists of 58 sequences derived from multiple published viral genomes. Given that the initial viral RNA sequences exceed 10000 nucleotides, we segmented them into modular structures of approximately 200 nucleotides each. The full curation process for these datasets is described in the Methods section. All test sets present unique challenges (bias, protein-assisted folding, structure model errors), making it difficult to establish a definitive ‘gold standard’ structure due to the sensitivity of RNA folding to experimental conditions and the presence of alternative structures. However, collectively, these test sets offer valuable insights for assessing variations in performance, particularly the capability of machine learning models to align with structure models derived from chemical probing data.

We evaluated four widely used algorithms on these curated test sets: RNAstructure Fold, EternaFold, MXFold2, and UFold, by calculating the F1 score distribution between the predicted and published structure models (Figure 1b). The F1 score, a commonly used metric in classification, balances precision and sensitivity in detecting correct base pairs, where a score of 1.0 signifies identical structures. All algorithms exhibited remarkable efficacy on the PDB dataset, typically achieving an F1 score around 0.9 (Figure 1b, Table 1). However, their performances notably declined on the more challenging test sets. For the viral structures, the average F1 score was around 0.7, and it was even lower for the long ncRNAs, with an average score of approximately 0.45 for RNAstructure Fold, EternaFold, and MXFold2 (Figure 1b, Table 1). Among these, UFold demonstrated the weakest performance, especially in predicting base pairs in long non-coding RNAs. This significant variation in performance across different test sets suggests that ML-based algorithms are primarily developed or trained for RNA types frequently encountered in the PDB and ArchiveII datasets, with limited exposure to newer RNA domains. It is notable that nearly half of the sequences in the ArchiveII test set exhibit more than 80% similarity (Supplementary Figure 1c). UFold’s lack of generalization is particularly pronounced when compared to MXFold2, a hybrid algorithm, as UFold is a fully end-to-end algorithm without built-in assumptions on base-pairing rules or RNA thermodynamics. To mitigate these limitations, we compiled a new dataset aimed at training ML-based algorithms to improve their adaptability and performance across diverse RNA domains.

**Table 1:** Performance of state-of-the art algorithms on diverse RNA types. Average precision, recall and F1 score between the predicted structure model of each algorithm against a published structure model across four distinct test sets: PDB, ArchiveII, Viral mRNAs, and Long ncRNAs. The cells are colored, with yellow corresponding to the maximum F1 score of 1, and dark blue to the minimum F1 Score of 0. The best model per test (column) is in bold.

| dataset | PDB |  |  | Archivell |  |  | Viral mRNA |  |  | Long ncRNA |  |  |
| --- | --- | --- | --- | --- | --- | --- | --- | --- | --- | --- | --- | --- |
|  | Precision | Recall | F1 | Precision | Recall | F1 | Precision | Recall | F1 | Precision | Recall | F1 |
| model |  |  |  |  |  |  |  |  |  |  |  |  |
| RNAstructure Fold | 0.90 | 0.91 | 0.89 | 0.52 | 0.59 | 0.55 | 0.69 | 0.74 | 0.71 | **0.46** | **0.52** | **0.49** |
| EternaFold | 0.88 | 0.91 | 0.88 | 0.56 | 0.65 | 0.60 | **0.75** | **0.81** | **0.77** | 0.45 | 0.47 | 0.46 |
| MXFold2 | **0.91** | 0.93 | **0.90** | 0.69 | 0.75 | 0.71 | 0.70 | 0.72 | 0.71 | 0.41 | 0.43 | 0.42 |
| UFold | 0.81 | **0.96** | 0.87 | **0.76** | **0.80** | **0.77** | 0.58 | 0.59 | 0.58 | 0.22 | 0.14 | 0.16 |

### A new database of biologically relevant RNAs to bridge the generalization gap

To address the generalization gap in RNA secondary structure prediction, we used DMS-MaPseq [22.] to probe 4,550 sequences from the 3’ end of human mRNAs and 1292 primary microRNA (pri-miRNA) sequences, which include the precursor hairpin and flanking regions (Methonds). Altogether, the distribution of lengths ranges from 200 to 1000 nucleotides (Figure 2a). Previous studies have shown that adding chemical probing constraints to RNAStructure yields models that are 90–100% identical to models obtained through crystallography, cryoEM, or phylogenetic covariation.[25., 26.]. To ensure high quality DMS signal we used a stringent coverage cut off of at least 3000 reads per base (Methods). The median signal for the DMS reactive bases, adenine and cytosine (AC), over uracil and guanine (GU) was 5.3 fold, with a two phase distribution between reactive and non reactive bases (Figure 2b). DMS reactivity for each RNA was then used as constraints in RNAstructure’s Fold algorithm to model the secondary structure. Even though the constraint parameters have undergone optimization [26], it is important to note that models derived from these data might not always align with the actual data. This discrepancy occurs because RNAstructure Fold produces an output regardless of its ability to find a suitable match for the data, a point further explored in the discussion on AUROC below.

**Figure 2:**
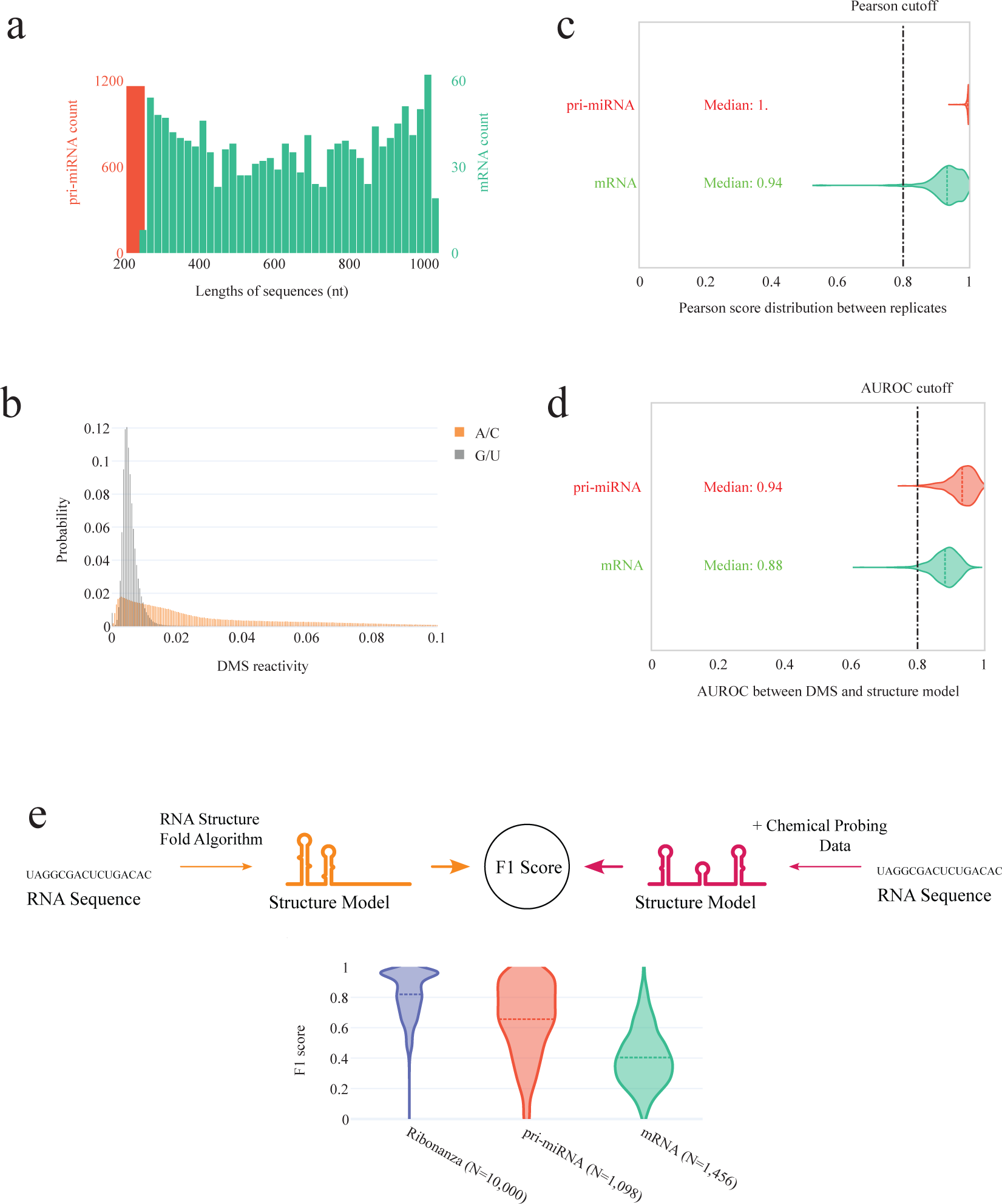
Data quality and complexity of RNAndria database. **(a)** Sequence length distribution for the pri-miRNA dataset (left scale,red) and mRNA dataset (right scale, teal). **(b)** Distribution of non-normalized DMS reactivity for the pri-miRNA and mRNA datasets. The non-reactive G/U bases are highlighted in gray with a mean reactivity of 0.006, while the reactive A/C bases are in orange with a mean reactivity of 0.03.**(c)** Distribution of Pearson correlation scores between DMS signal replicates. The dotted line corresponds to the threshold of 0.8 used for filtering. **(d)** Distribution of AUROC between the structure models and DMS signals. The dotted line corresponds to the threshold of 0.8 used for filtering. **(e)** Distribution of F1 score between structures predicted by RNAstructure Fold from sequence alone compared to RNAstructure Fold with chemical probing constraints, for Ribonanza, Pri-miRNA, and human mRNA regions (mRNA) datatest.

We assessed the quality of the final dataset in several ways. Over 70% of sequences underwent dual independent probing. The correlation between the DMS signal replicates is depicted in Figure 2c. The reproducibility of the signals is highlighted by a high mean Pearson correlation between DMS replicates of 0.95 for the mRNA and 0.99 for the pri-miRNA. To evaluate the agreement of the derived structure models with the DMS data, we utilized the Area Under the Receiver Operating Characteristic curve (AUROC). The ROC curve plots the trade-off between precision (true positive rate) and sensitivity (false positive rate) of a binary classifier, as it differentiates between paired and unpaired bases at varying thresholds of DMS signal intensity. At each threshold, it computes the true positive rate and false positive rate in comparison to a given structure model. An AUROC score of 1.0 signifies the existence of a specific threshold where all DMS-reactive positions correctly align with loop regions and all DMS-unreactive positions with stem regions, demonstrating perfect agreement with the observed data. Although a high AUROC does not validate the model (there could exist two or more conformations that perfectly fit the data), a low AUROC indicates the model is incorrect, either due the presence of alternative structures or poor data quality. Indeed, “gold standard” structures determined by crystallography or NMR have high AUROCs (>0.8) [27.]. We present the AUROC values for each structure in Figure 2c and our newly established database, RNAndria, and exclude any structure with an AUROC below 0.8 (Methods).

We highlight the importance of integrating DMS signals into our structure models by comparing the performance of RNAstructure Fold with and without DMS constraints. Our analysis reveals a significant variation in model performance for mRNA and pri-miRNA datasets—F1 scores range from 0.3 to 0.7—when relying solely on sequence data versus when DMS constraints are included (Figure 2d). This discrepancy in F1 scores aligns with the marked improvements in RNAstructure Fold’s accuracy for ribosomal RNA, which jumps from approximately 50% to 90% with the incorporation of chemical probing data [25]. The notably low F1 scores for RNAstructure Fold on the RNAndria dataset indicate the challenge of predicting complex structures in the absence of probing data. In contrast, models derived from Ribonanza [17], the only other extensive chemical probing dataset, show minimal variance with or without chemical constraints (Figure 2d). This suggests that the sequences in Ribonanza fold into motifs that are more predictable. The stark difference highlights the distinct value of our dataset in enhancing model training, enabling better generalization across a broad spectrum of RNA structural motifs.

RNAndria post filtering for high read depth includes 1,456 mRNA and 1,098 pri-miRNA structures. This dataset is available in an online database-https://rnandria.org/, which allows users to search and filter by data features including species, condition, dataset, and more. Researchers can then visualize and download their data of interest. We included visualization of both the secondary structure and DMS signal (Figure 3). The secondary structures, along with their DMS reactivity per base, was performed using RNArtist (Methods), a tool suited for representing long and complex RNA structures.

**Figure 3:**
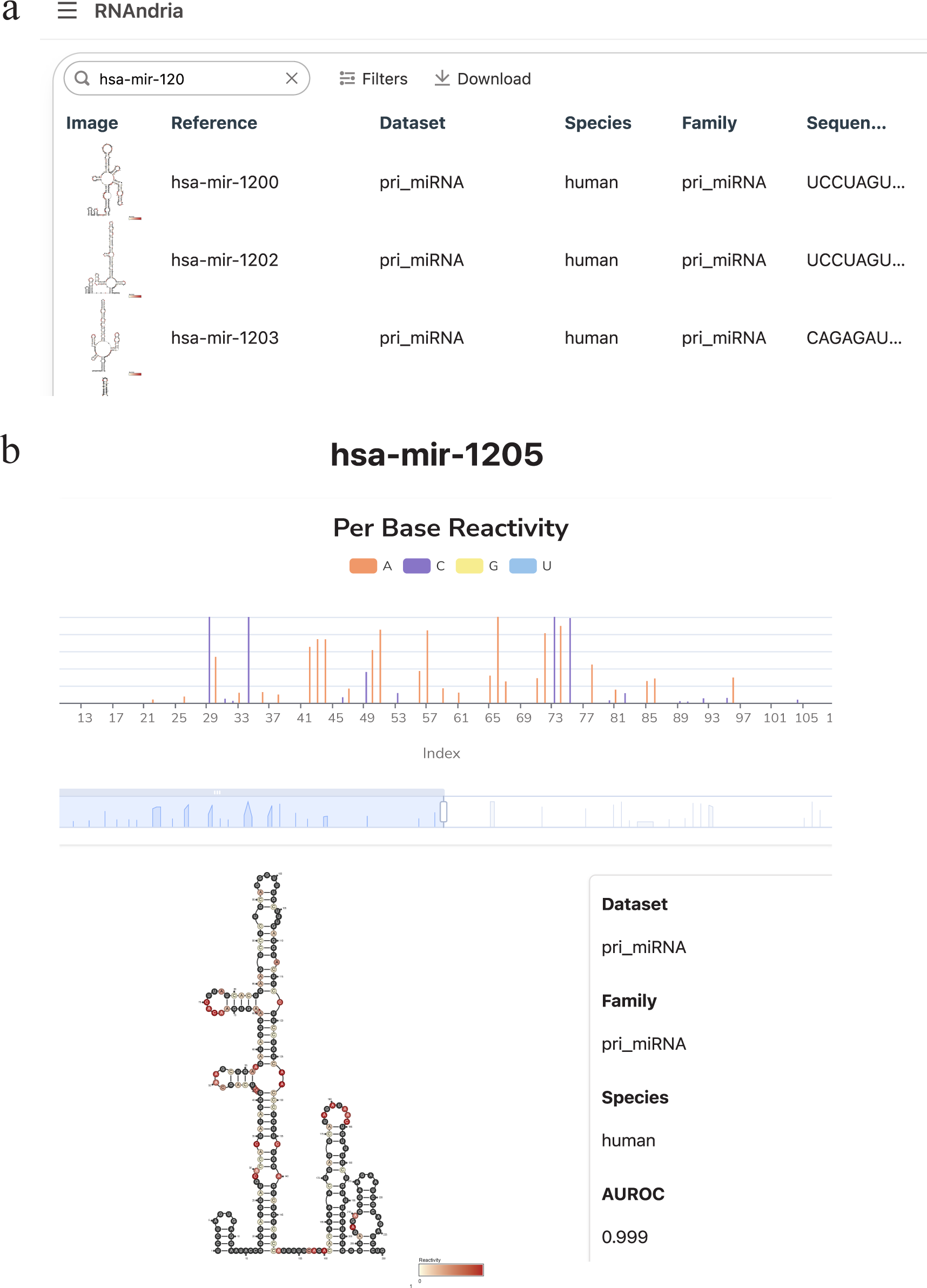
RNAndria database architecture. **(a)** Main page of the online database RNAndria, showing the browsing, filtering, searching and download features.**(b)** Sub page example for one sequence of the pri-miRNA hsa-mir-1205, showing its secondary structure, DMS reactivity, and metadata.

### A novel architecture for secondary structure prediction

We next explored a new neural network architecture for RNA structure prediction adapted from the Evoformer module of Alphafold [23.]. This model, which we called eFold, consists of four blocks depicted in Figure 4. The core design of each eFold block contains two channels: one channel processes the sequence representation through self-attention layers, and the other handles pairwise representation via ResNet convolutional layers. Specialized connections between layers in each channel facilitate effective information exchange. Convolutional neural networks (CNNs) have been heavily used in many deep learning structure prediction models, and are thus integral to our architecture. Their suitability for handling image-like representations of pairing matrices, along with their computational efficiency, makes them preferable over the graph neural network blocks found in the original Evoformer. However, CNNs typically have limitations in capturing long-range interactions, an issue we address by incorporating self-attention layers. These layers process the sequence representation without range limitations, effectively complementing the CNNs.

**Figure 4:**
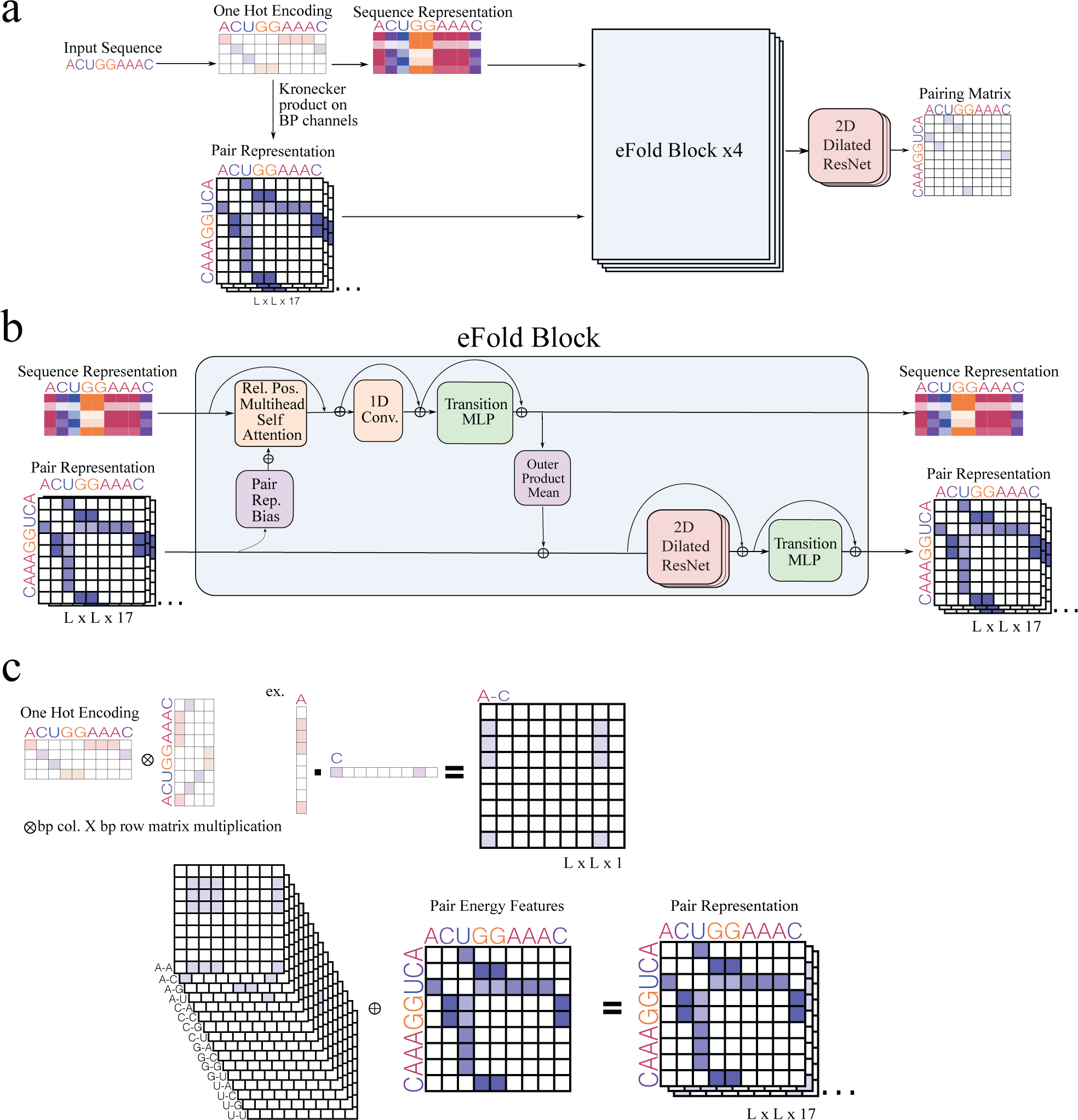
eFold architecture. **(a)** High level schematic of the eFold architecture, with the two input representations and the output pairing matrix. **(b)** Low level view of one eFold block, showing the sequence and pair representation channels, as well as the inter channel connections. **(c)** Details on the computation of the input pair representation from sequence. Similar to UFold, we use 16 binary matrices to represent all base interactions, and one channel to represent the energy of A-U, G-C and G-U pairs.

As shown in Figure 4a, the sequence representation is initialized using a simple table embedding, while the pair representation adopts a method similar to that in the UFold architecture [7.]. This setup encodes the sequence as a binary map representing all 16 possible base pair interactions and includes an additional channel that factors in the energy of different pairs. Following the eFold blocks, only the pair representation is advanced further. It is then processed through a ResNet block, resulting in the generation of the pairing probability matrix. This novel architecture is designed to balance the strengths of both CNNs and self-attention layers, aiming to enhance the accuracy and efficiency of RNA secondary structure prediction from sequence data.

### Enhanced performance of eFold in predicting the structure of long RNAs

Our model underwent a two-phase training process. Initially, we utilized existing databases, including bpRNA and Ribonanza, to compile a comprehensive pre-training dataset comprising over 120,000 unique sequences and structures. To ensure data quality, redundant sequences were filtered out using BLAST [28.], and low-quality structure models were discarded (Supplementary Figure 2b). Additionally, we incorporated 220,000 sequences from RNAcentral and employed RNAstructure Fold to generate their structures. This synthetic dataset facilitated our model’s learning of RNAstructure Fold’s basic pairing rules, enabling better generalization across new families and sequence lengths. A summary of the datasets used for pretraining, finetuning and testing is presented in Table 2. The pre-training phase alone permitted our model to surpass the performance of the comparable end-to-end model, UFold, on the long non-coding RNA test set, achieving a mean F1 score of 0.4 versus UFold’s 0.16 (Figure 5a). While performance on other test sets was similar, an exception was noted with ArchiveII, where UFold exhibited higher accuracy (Table 3). Limiting the pre-training to only the bpRNA or RNAstralign dataset resulted in increased accuracy on ArchiveII but a failure to generalize effectively across the other test sets, akin to UFold’s performance (Supplementary Figure 3a), highlighting the inherent bias in datasets selected for pre-training. In the subsequent phase, we fine-tuned our model with curated mRNA and pri-miRNA datasets, which led to improved mean F1 scores across all test sets, demonstrating the importance of new and diverse data in enhancing model generalization (Figure 5a, Table 3). The final eFold model outperformed in the two most challenging test sets, registering a mean F1 score of 0.73 on the viral mRNA compared to UFold’s 0.58, and 0.45 on the long ncRNA, against UFold’s 0.16. For an illustrative example, we present a comparison of a functional element from the lncRNA PAN structure model of Kaposi’s sarcoma herpesvirus (1.1kb), as predicted by various algorithms, against the reference structure derived from ex-vivo deprotonated RNA probed with SHAPE (Figure 5b) [29.].

**Figure 5:**
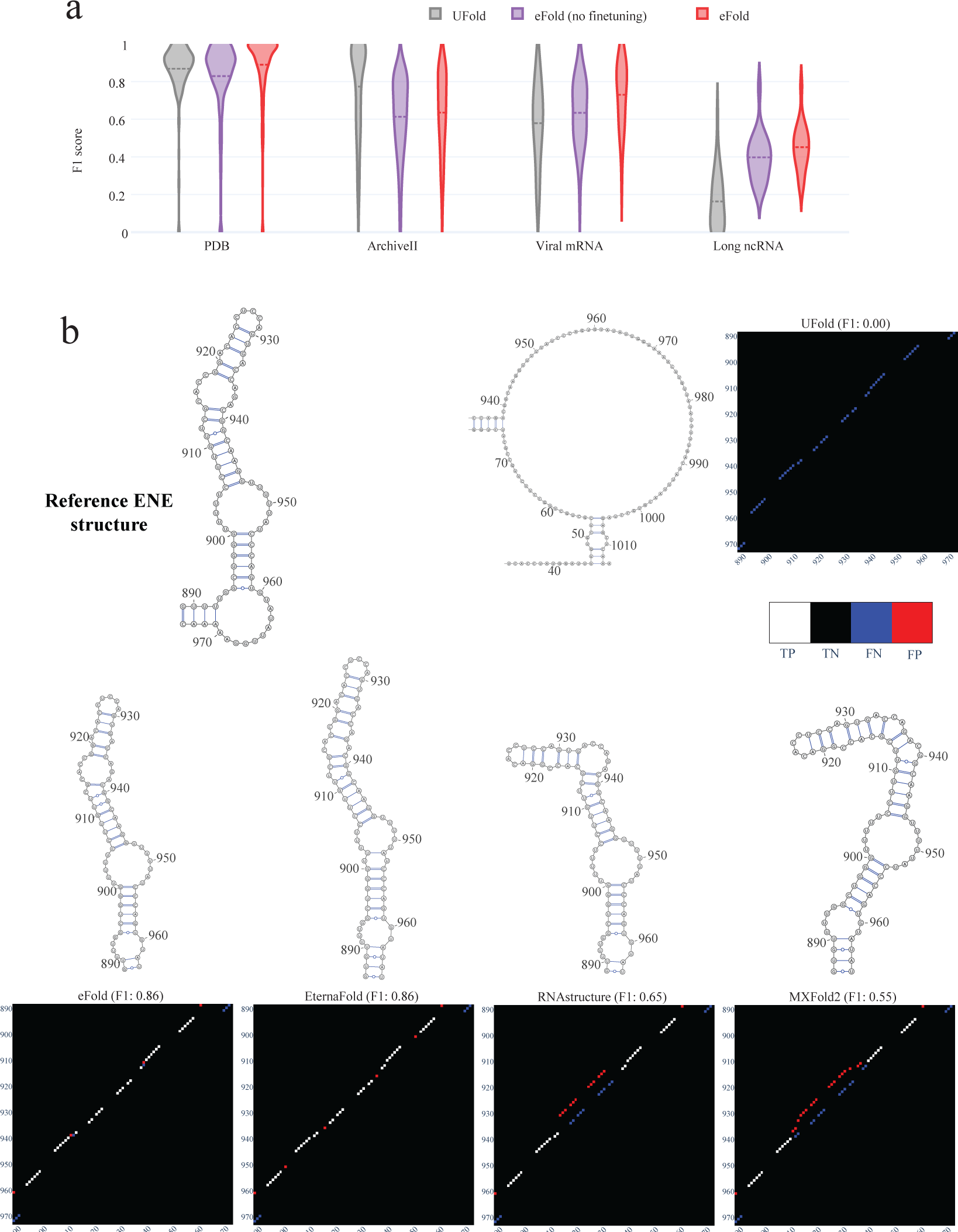
eFold performance. **(a)** F1 score distribution between each model prediction and the ground truth, for the four test sets. In purple is eFold only pre trained on bpRNA, Ribonanza and the synthetic dataset, while eFold in red is pre trained then fine-tuned on the mRNA and pri-miRNA datasets. **(b)** Secondary structure model of the ENE (Element for Nuclear Expression), a functional element within Polyadenylated Nuclear RNA (PAN) of Kaposi’s sarcoma herpesvirus. The full secondary structure is predicted with each algorithm, then the ENE is drawn using VARNA, and the predicted and reference base pairing matrix are plotted next to each model, showing the correct base pairs in white (True Positive, TP), the missing base pairs in blue (False Negative, FN) and incorrect base pair predictions in red (False Positive, FP).

**Table 2:**
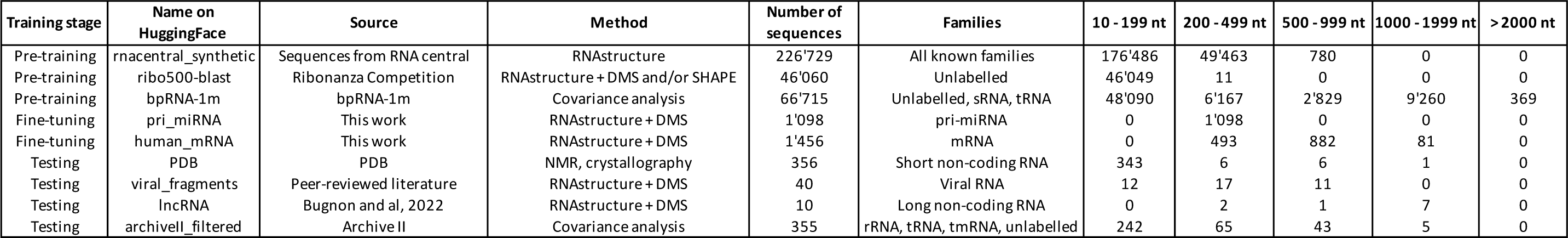
Summary of training datasets. Description of each dataset used either as pretraining, fine-tuning or testing, along with their curation method, source of data, family and length information.

**Table 3:**
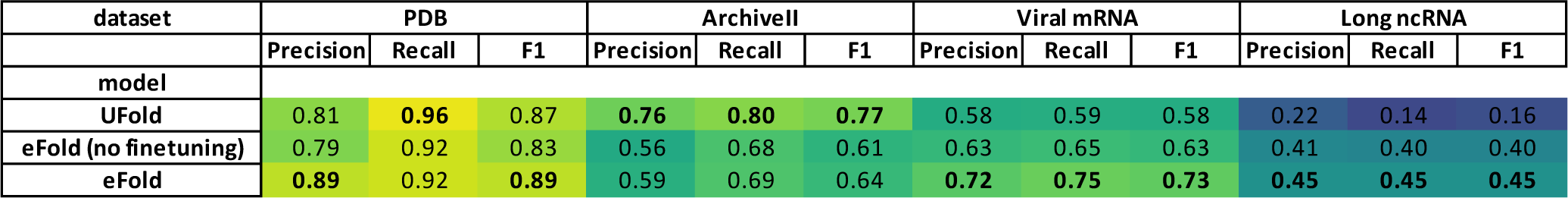
Performance of end-to-end models. Average precision, recall and F1 score between the predicted structure model of UFold, eFold without fine-tuning and eFold with fine-tuning against a published structure model, across four distinct test sets: PDB, ArchiveII, Viral mRNAs, and Long ncRNAs. The cells are colored, with yellow corresponding to the maximum F1 score of 1, and dark blue to the minimum F1 Score of 0. The best model per test (column) is in bold.

The contributions of individual training components to eFold’s performance were examined through ablation tests (Supplementary Figure 3b). In comparing the architectures of UFold and eFold, each trained on the identical pre-training dataset, we noted the most pronounced improvement in the long non-coding RNA dataset and a modest enhancement in viral structure predictions. This significant performance boost, especially for long RNA sequences, can likely be attributed to eFold’s architecture, which facilitates both long and short-range interactions across the network. Additionally, the implementation of relative positional embeddings plays a crucial role in enabling the model to generalize to sequences of varying lengths. The fine-tuning stage, particularly with the RNAndria dataset, marked the next most significant improvement, affecting both viral and long non-coding RNA structures. We conducted experiments with fine-tuning using exclusively the mRNA or pri-miRNA datasets. The findings suggested that the mRNA dataset was more pivotal for achieving broad generalization. Performance closely matched the baseline when training was confined solely to the mRNA dataset (Supplementary Figure 3b), likely due to its larger size and the diversity in sequence lengths it offers, which are crucial for generalizing to the long non-coding RNA test set.

## Discussion

In this study, we addressed two significant challenges in the field of RNA secondary structure prediction: the limited generalizability of existing algorithms and the necessity for a robust, end-to-end predictive model. We highlighted the difficulty in predicting the structures of a broad class of biologically relevant RNA molecules, including mRNAs, viral RNAs, long ncRNAs, and pri-miRNAs, with conventional algorithms. By curating a list of published structure models obtained through chemical probing, we demonstrated the shortcomings of machine-learning-based approaches in generalizing to novel RNA types beyond their training sets. Our work was directed along two paths: first, the compilation of a new, diverse dataset of RNA structures; and second, the development of eFold, a cutting-edge model for RNA secondary structure prediction.

The dataset we created encompasses secondary structures for 1,456 human mRNA regions and 1,098 pri-miRNAs, as determined through chemical probing. This collection significantly broadens the spectrum of RNA structures available for research, addressing a vital deficiency in the current data landscape. The prevailing datasets, predominantly composed of short ncRNAs, have not sufficiently supported the development of algorithms capable of accurately predicting RNA structures. By integrating a wide range of RNA families and lengths, our objective was to enhance the predictive models’ ability to generalize.

eFold, drawing inspiration from the Evoformer block of AlphaFold and incorporating elements of traditional architectures, marks a notable advancement in the prediction of RNA secondary structures. This end-to-end deep learning model was trained using the RNAndria database and over 300,000 secondary structures from diverse sources. Our evaluations reveal that eFold significantly outperforms existing state-of-the-art end-to-end methods, especially in predicting complex structures, such as those of long ncRNAs. This achievement highlights the critical importance of diversifying the training dataset in terms of structure diversity, length, and complexity to augment the precision of ML-based models.

This research underlines the pressing need for datasets that encompass a broad array of long RNA sequences and complex structures. At present, the accuracy of predictions for a curated selection of published long ncRNA structures ranges from 40% to 50%. This stands in stark contrast to the 90-100% accuracy rates achieved for short ncRNAs within the PDB. The introduction of the eFold model and the RNAndria database represents a significant stride towards mitigating this disparity. With the inclusion of a larger and more varied dataset, potentially encompassing alternative structures, achieving a level of predictive precision for long RNAs comparable to the high accuracy rates currently observed for short ncRNAs becomes a tangible goal. Such precision is essential for unraveling the complex roles and functions of RNA structures in biological systems.

## Acknowledgment

We thank Prof. Dan Herschlag for providing valuable feedback and helping us improve this work. This work was supported by the Burroughs Wellcome Fund G-1018246.01

## Author Contributions

A.L,Y.M., and S.R. conceived and designed the project with input from C.K. A.L. and Y.M. build and trained eFold with input from C.K. and A.K. F.W, J.A. carried out experiments and design of the sequencing and structure probing approach. M.A. designed the human mRNA dataset.

D.S performed DMS-MaPseq for the mRNA sequences. C.LK. built the web interface. F.J. drew all the structure models in the RNAndria database.

## Declaration of Interests

The authors declare no competing interests

## Data availability

All processed datasets are accessible publicly on Huggingface: https://huggingface.co/rouskinlab. The processed pri-miRNA and mRNA datasets are available at rnandria.org and the raw fastq files are accessible on GEO (GSE262014)

## Methods

### Library design

To generate the human mRNA library, all human protein-coding transcripts were downloaded from GENCODE Release 43 [30.]. Every transcript with a non-AUG start codon or without an annotated 5’ UTR, CDS, and 3’ UTR was discarded. To focus on 3’ UTRs and maximize the likelihood of successful RT-PCRs, for each gene, the longest common suffix (i.e. subsequence beginning at the 3’ end) among all isoforms was selected. Each gene was discarded if it had no annotated name, if its longest common suffix was shorter than 253 nt, or (to minimize mispriming and misalignment) if its longest common suffix contained any 41-mers that were present in the longest common suffix of any other gene. To verify that alignment to the wrong gene would be minimal, a simulated FASTQ file containing every 128 nt segment in 64 nt increments from every gene’s longest common suffix was aligned to all longest common suffixes with Bowtie2 v2.5.1 [31.].

Primers for human mRNAs were designed against the longest common suffixes of all remaining genes using Primer3 v2.6.1 [32.] with the parameters PRIMER_TASK=generic, PRIMER_PICK_LEFT_PRIMER=1, PRIMER_PICK_INTERNAL_OLIGO=0, PRIMER_PICK_RIGHT_PRIMER=1, PRIMER_PRODUCT_SIZE_RANGE=253-1021, PRIMER_OPT_SIZE=28, PRIMER_MIN_SIZE=20, PRIMER_MAX_SIZE=36, PRIMER_WT_SIZE_LT=0.0, PRIMER_WT_SIZE_GT=0.0, PRIMER_MIN_TM=65.0, PRIMER_OPT_TM=75.0, PRIMER_MAX_TM=80.0, PRIMER_PAIR_MAX_DIFF_TM=10.0, PRIMER_GC_CLAMP=0, PRIMER_MAX_NS_ACCEPTED=0, P3_FILE_FLAG=0, PRIMER_EXPLAIN_FLAG=1, SEQUENCE_OVERHANG_LEFT=TAATACGACTCACTATAGGG. A T7 promoter ending in three G bases was prepended to every forward primer. Genes that received no primer pairs were discarded; for those that received more than one pair, the pair that would produce the longest amplicon was selected. The sequences of the amplicons (including only the last 3 Gs of the T7 promoter) were used as the reference sequences in the subsequent analysis.

To select amplicons for verification in HEK 293T cells, four microarray datasets of gene expression in HEK 293T cells from two studies [33.] were downloaded from the National Center for Biotechnology Information’s Gene Expression Omnibus [34.] (accession numbers GSM389756, GSM389757, GSM609202, and GSM609205). Microarray probe IDs were converted to gene names using DAVID [35.]. The amplicons of at most 600 nt (the maximum paired-end read length on Illumina sequencers) were ranked by the minimum expression level of their genes among the four datasets (to be conservative); the 96 highest-ranked amplicons were chosen for mutational profiling in HEK 293T cells.

To generate the pri-miRNA library, human genome annotations were downloaded from miRBase[36.]. After removing redundant sequences, 1292 miRBase hairpins were selected for downstream library creation. The miRBase hairpins were then padded with their flanking genomic sequence to a total of 192 nt. These 1292 sequences were then separated into 3 sub-libraries to reduce jackpotting during library processing. To each construct, a sub-library specific pair of 19 nt universal primer binding regions were added to minimize primer associated PCR and reverse transcription bias, bringing the full length library size to 230 nt. The final sequences were then synthesized as an oligo pool (Agilent Technologies).

### DMS data generation

For the in vitro transcription of the human mRNA library, all the designed amplicons were amplified via PCR, adding a T7 promoter sequence at the forward primer (5′ TAATACGACTCACTATAG 3′) using a 2X PCR PreMix (Syd Labs, Cat. MB067-EQ2N), human cDNA (ZYAGEN, Cat. HD-UR-40) and an INTEGRA VIAFLO384 channel pipette. For each 384 plate, 5 µl of each individual reaction were pooled together and purified using a Zymo DNA clean up kit (Cat. D4003). The PCR product pool was used as template for the in vitro transcription, using the T7 MEGAscript kit (Thermo Fisher, Cat. AMB13345) according to the manufacturer’s instructions. After DNase treatment, the transcribed RNAs were purified using a Zymo RNA clean up kit (Cat. R1017). 10 µg of the purified RNAs were diluted in 10 µl of nuclease free water, denatured for 1 minute at 95 degrees and placed on ice for 2 minutes. 88.5 µl of the refolding buffer (sodium cacodylate 397.5 mM and MgCl_2_ 6 mM final) were added and the RNAs were left refolding at 20 minutes at 37°C. After, 1.5 µl of DMS (dimethyl sulfate) was added to obtain a 1.5% final concentration, for 5 minutes at 37°C, while shaking at 600 RPM. The reaction was stopped by adding 60 µl of β-mercaptoethanol, followed by Zymo RNA clean up.

Libraries were generated as previously described [37.], and the indexed samples were sequenced in a Illumina NextSeq 2000.

For the pri-miRNA library, *in vitro* transcription templates were prepared via 8 cycles of PCR using 2x Q5 Master Mix (New England Biolabs, Cat. M0492S) with a T7 promoter containing forward primer as previously described. Following column clean up (Zymo Research, Cat. D4013), 100 ng of template DNA was used in a 2 hour *in vitro* transcription reaction using the HiScribe T7 High Yield RNA (New England Biolabs, Cat. E2040S) kit according to the manufacturer’s instructions. Following DNase digestion (Invitrogen, Cat. AM2238) and column clean up (Zymo Research, Cat. R1017), 1 ug of RNA from each library was DMS treated separately as described above, substituting a 1.5% final DMS concentration for 2.5%. Following DMS treatment, the samples were reverse transcribed with Induro Reverse Transcriptase (New England Biolabs, Cat. M0681L) for 30 minutes at 55°C using the appropriate reverse primer for each library according to the manufacturer protocol. After alkaline hydrolysis with 1 ul of 4M NaOH, cDNA was cleaned and concentrated using Zymo Oligo Clean and Concentrator (Zymo Research, D4060). Sub-library specific primers were then used to amplify 1 ul of cDNA template for 24 cycles using Advantage 2 PCR Mix (Takara Bio, Cat. 639206). Finally, library indexing was carried out with the NEBNext Ultra II DNA Library Prep Kit for Illumina (New England Biolabs, Cat. E7645S) and sequenced on a NextSeq 1000 P2 cartridge (Illumina, Cat. 20046813) in accordance with the manufacturer’s instructions.

### DMS data processing and filtering

For the processing of raw reads, the fastq files were handled using SEISMIC-RNA [38.], which aligned the reads to the reference sequence and quantified mutations per base. Sequences with less than 3000 aligned reads or where less than 50% of bases had coverage of at least 3000 reads were excluded. In some cases, an untreated RNA sample was sequenced to provide a baseline, aiding in the identification of abnormally high mutation rates (>0.3) that might indicate inaccuracies in the reference sequence. If mutations consistently occurred towards a specific base, the reference sequence was updated; otherwise, the sequence was discarded. Biological replicates were analyzed to check experimental consistency. Replicates with a pearson correlation below 0.8 were removed, while those with higher correlation were averaged for further use. Bases were masked with a value of −1000 if their coverage was below 3000, if they were G or U bases, or if they were part of the primer region in the pri-miRNA dataset. Signal normalization was then performed per reference, using the 95th percentile as the maximum value, with values above this threshold clipped to 1. The RNAstructure Fold algorithm, incorporating the DMS signal as a constraint, was then applied. Structures with an AUROC of less than 0.8 in relation to the DMS signal were excluded. Through these measures, 15% of the pri-miRNA dataset and 70% of the human mRNA dataset were removed, ensuring the high quality of the remaining data.

### Data quality assessment

Pri-miRNA has replicates for 99% of the references and human mRNA has replicates for 50% of the references. This is because we have very stringent filtering steps, so some references were valid in one experiment and not in another. We estimated the reproducibility of our data by comparing our replicates. The DMS signals replicates were compared using the Pearson correlation score. The median Pearson score was 1.0 for pri-miRNA and 0.94 for mRNA. Note that in the final dataset, both replicates were aggregated into a single signal.

We also estimated the robustness of the data as the number of reads decreases. The DMS-MaPseq method consists in counting the mutated bases at each position across a large number of reads. To estimate the error due to the number of reads, we model the mutation fraction as a binomial distribution divided by the number of trials. We bootstrap 10 new dms signals for each reference from the original signal using this model. We predict the structure using RNAstructure and these DMS signals. The structures are systematically the same when predicted with the original signal and the bootstrapped signals, suggesting that subsampling error doesn’t have an influence on the structure prediction.

### Test sets

For our test sets, we sourced data from a variety of published papers by different research groups. In selecting entries from the Protein Data Bank (PDB), we focused on those classified under “Polymer Composition” as RNA, and with a “Number of Assemblies”, “Number of Distinct Molecular Entities”, and “Total Number of Polymer Instances” all set at 1. This approach yielded 356 entries, which were then converted from tertiary to secondary structures using the RNApdbee webserver, applying the default settings: 3DNA/DSSR as the conversion software and the hybrid algorithm method.

The ArchiveII dataset was downloaded from Mathew lab website (https://rna.urmc.rochester.edu/publications.html). We only removed sequences from ArchiveII that were also present in bpRNA and RNAstralign, as these are commonly used train sets. We call this subset “ArchiveII_filtered”

The viral structures dataset was created by segmenting long viral RNA sequences into smaller, independent modules. We utilized the HIV structure from Watts et al. [39.], the SARS-CoV structure from Lan et al. [40.], the Hepatitis virus structure from Mauger et al.[41.], and the Alphavirus structure from Kutchko et al. [42.]. For each of these structures, we identified modular structures characterized by fully closed loops and high agreement between the structure and chemical probing data (AUROC > 0.8). After segmenting the chemical probing signal (DMS or SHAPE), RNAstructure was rerun with each fragment. Fragments were retained if the new structure aligned with the corresponding segment of the chemical probing signal (AUROC > 0.8 and F1>0.8).

The long non-coding RNA (lncRNA) dataset was sourced from Bugnon et al. [11.], with the only modification being to cut sequences exceeding 2,000 nucleotides in length, using the same method as for the viral structures. We didn’t use the last filtering step of the cutting process as we found it made almost no difference in the test results of all algorithms and models.

### External databases curation

We combined the public databases bpRNA [13.] and Ribonanza [17.] into a pre-training dataset. We applied the following filtering steps: first, we removed duplicate sequences within the databases. We only kept sequences with the canonical bases ACGU. The T bases were converted to U. We filtered out the sequences below 10 nucleotides. We removed sequences for which we have a sequence but no structure. We removed sequences that are common between datasets. For ribonanza specifically, we applied the following filtering: we filtered low-reads and low S/N ratio data. The cutoff was set to more than 500 reads and a S/N ratio - a quality indicator provided by the Ribonanza dataset - greater than 1. The data includes structures predicted with EternaFold. To ensure that the EternaFold-predicted structure was matching the signal, we computed the AUROC between the structure and the one of the chemical probing signals for each sequence. We used DMS by default, and SHAPE if DMS was filtered out. If the AUROC was below 0.8, the structure was filtered out. We removed redundant sequences within the dataset using BLAST. The primers were masked. If two sequences had over 80% matches on a sequence of over 112 nucleotides, we kept only the best covered sequence. Note that 112 nucleotides correspond to 80% of the most represented length minus the primers.

The last dataset used for pretraining is a synthetic dataset. We gathered sequences from RNACentral by sampling uniformly from all RNA clans, then added a balanced proportion of mRNA and viral sequences. The final dataset contains 227,000 sequences up to 512 in length. We predict the structure of each sequence using RNAstructure Fold

### Structure visualization

All the secondary structures available in RNAndria, along with their DMS reactivity per base, have been plotted from the command line using the tool RNArtistCore (https://github.com/fjossinet/RNArtistCore). Built with Kotlin (https://kotlinlang.org/), RNArtistCore follows an heuristic approach to identify a non-overlapping 2D layout for an RNA secondary structure.

At first, RNArtistCore computes the structural domains (helices and junctions) from paired positions. In 3D, junctions connect and orient helices in order to produce the biologically relevant 3D architectures. Each junction is associated to a class depending on the number of helices connected (apical loops, inner loops, 3-way junctions, 4-way junctions….).Starting from the 5’-end, RNArtistCore is processing each junction in order to find orientations for its helices in order to avoid overlapping between upstream and downstream structural domains in a 2D sketch.

Except for apical loops, the drawing engine implemented in RNArtistCore splits the connected helices in a junction between a single ingoing helix and outgoing ones. The modeling of RNA concepts of RNArtistCore links each junction class to a default positioning for its outgoing helices, relatively to the orientation of its ingoing one. If the default position for an outgoing helix produces an overlap between the next junction and at least one upstream structural domains, RNArtistCore searches for a new position among those still available. When the orientation of an outgoing helix has been defined, this helix becomes the ingoing one for the next junction to be processed.

Once a non-overlapping layout found, it is used to produce an SVG drawing according to some user-defined instructions. RNArtistCore provides a simple and expressive syntax (a.k.a « Domain Specific Language ») to allow the user to parameter the final result. The instructions are described in a script whose name and location has to be specified on the command line. Among all the options available (details levels, colors, line widths,..), one can easily link a dataset (like DMS reactivities) to the structure and map its values to a color gradient.

Some 2D RNA structures can contain nested junctions for which RNArtistCore could not be able to find any non-overlapping organization. In such cases, RNArtistCore can be configured to go backwards in order to test different layouts for upstream junctions that could unlock the nested ones. For this study, we allowed RNArtistCore to go back until 10 previous junctions if necessary.

### Models architecture

Model inputs are RNA sequences only. The output is a 2D RNA structure prediction in the form of a pairing matrix. The sequence is one-hot encoded and then fed through a learned embedding layer (dimension = 64). This learned representation is the input to the eFold module. The first step in this module is constructing the pair representation. The one-hot encoded sequence is transformed using a Kronecker outer product (specified here [3.]) to create an LxLx17 pair representation. The sequence representation goes through a relative positional multihead self-attention module with bias from the paired representation. This module consists of a multiheaded attention mechanism with learned relative position encodings [43.] and pair representation bias added to the logits prior to the softmax function. The sequence representation goes through a feed-forward layer followed by a 1-D convolution module. This includes pointwise convolutions followed by depthwise convolution (number of output channels = 128) and finally another pointwise convolution (number of output channels = 64). This is followed by another feedforward layer and a 2-layer deep sequence representation transition MLP. There are skip connections between each module. The output of the transition MLP is transformed using an outer mean product and added to the pair representation for the second trunk of the eFold module. The pair representation is sent through a dilated 2D resenet with 2 residual-blocks of depth. The output is followed by a 2-layer deep pair representation transition MLP. The eFold module is repeated four times prior to the output layers. Pairing matrix predictions are sent through a final 2D residual block head (output dimensions = LxLx1). Loss is calculated with binary cross entropy. The pairing matrix is post-processed by enforcing constraints such as no sharp loops and only one nucleotide can be paired with another. We reuse the same optimization method as UFold to find the pairing matrix respecting the constraints with minimum modification of the model output.

### Training and evaluation

The model is developed using the Pytorch and Lightning libraries. We use the Binary Cross Entropy (BCE) loss for structure prediction. We use a cluster of 8 Nvidia RTX4090 and train all models and experiments using the DDP strategy for 20 epoch. Because the results can vary from epoch to epoch, we average the weights of the last 5 epochs (epochs 16 to 20) to get the final weights used in the experiments. Due to large memory constraints, we run the model with a batch size of 1, and thus don’t require any padding. Using gradient accumulation, the effective batch size is 256. We train with Adam optimizer with a learning rate of 0.0003.

The test sets are run separately on one GPU to ensure maximum precision in the metric computations. All the code is freely available on Github at https://github.com/rouskinlab/efold and the datasets on Huggingface at https://huggingface.co/rouskinlab.

## Supplementary Figures

**Supplementary Figure 1:**
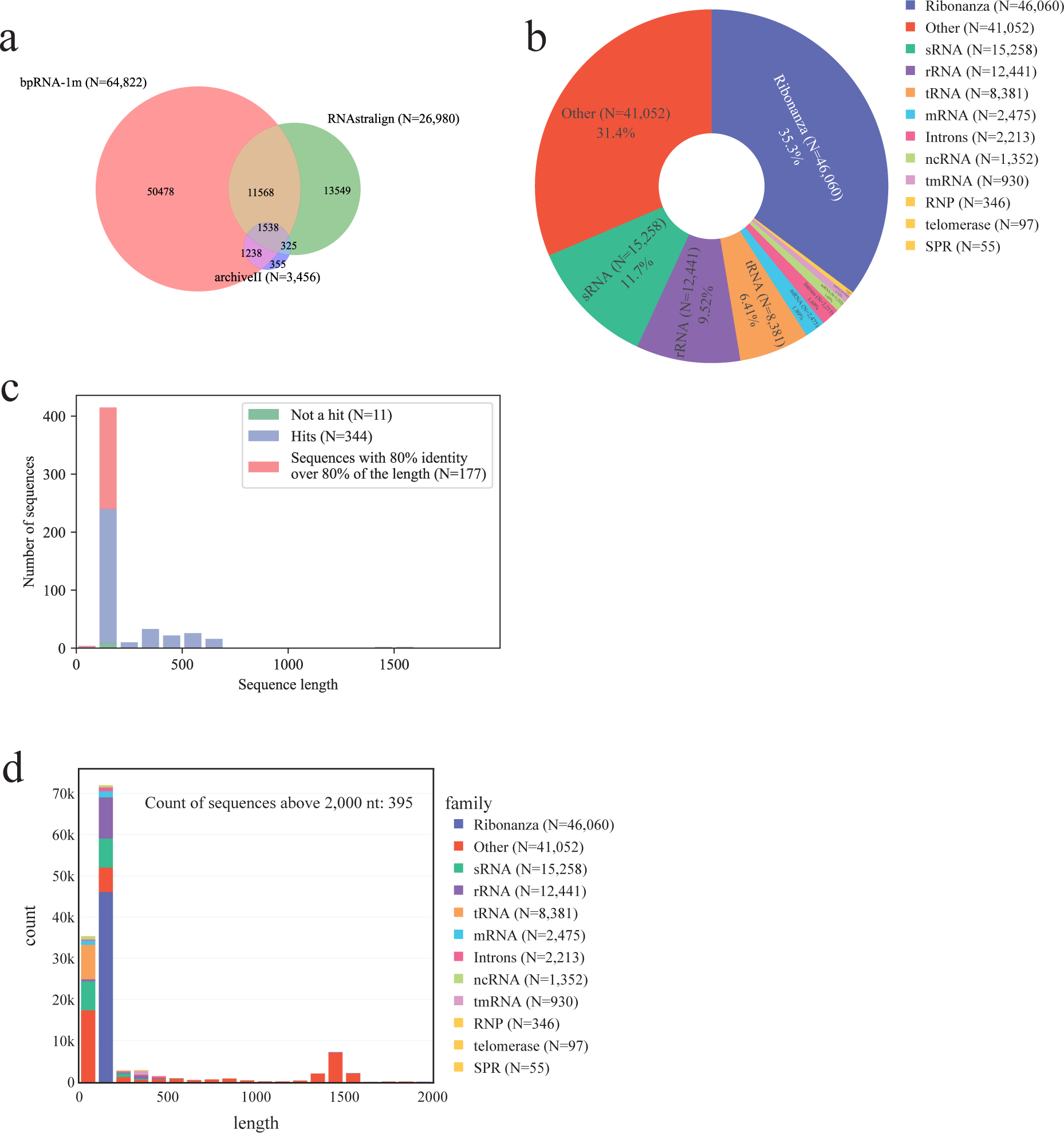
Representation of the external databases and of the families and lengths of the pre-train set. **(a)** The common sequences between bpRNA, RNAstralign and ArchiveII are represented using a Venn diagram. **(b)** The distribution in families of the compiled train set, displayed as a pie chart. Ribonanza counts as a family since we don’t have labels for it. **(c)** BLAST analysis of the ArchiveII test set compared to bpRNA and RNAstralign, after removing common sequences. **(d)** The distribution of the length of the compiled train set, displayed as a histogram.

**Supplementary Figure 2:**
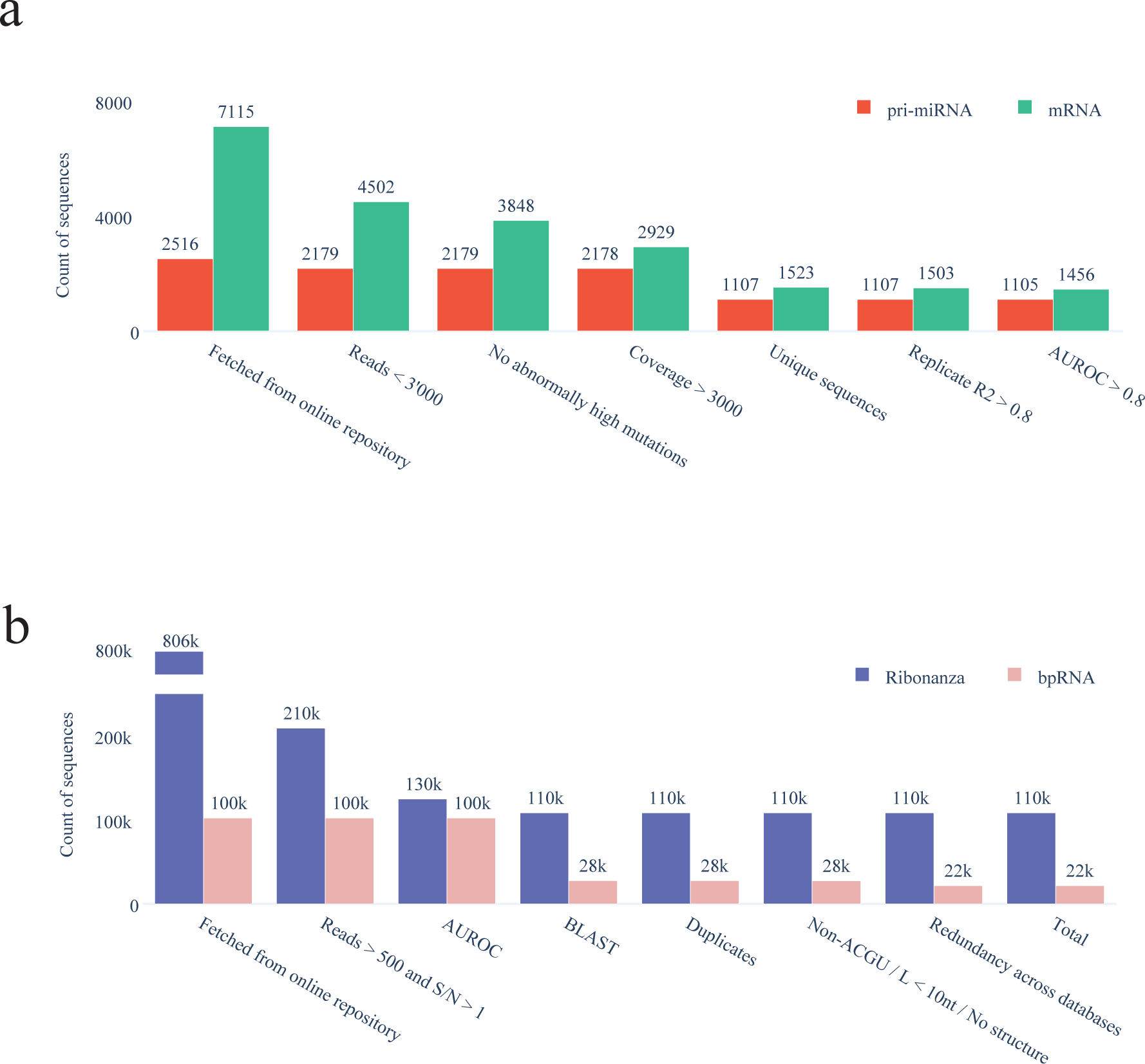
Filtering steps of the datasets. **(a)** Dataset size at each filtering step for the pri-miRNA (red) and mRNA (green) datasets before fine-tuning. **(b)** Dataset size at each filtering step of the bpRNA (pink) and Ribonanza (blue) datasets before pretraining. Note that the y axis is discontinuous for clarity.

**Supplementary Figure 3:**
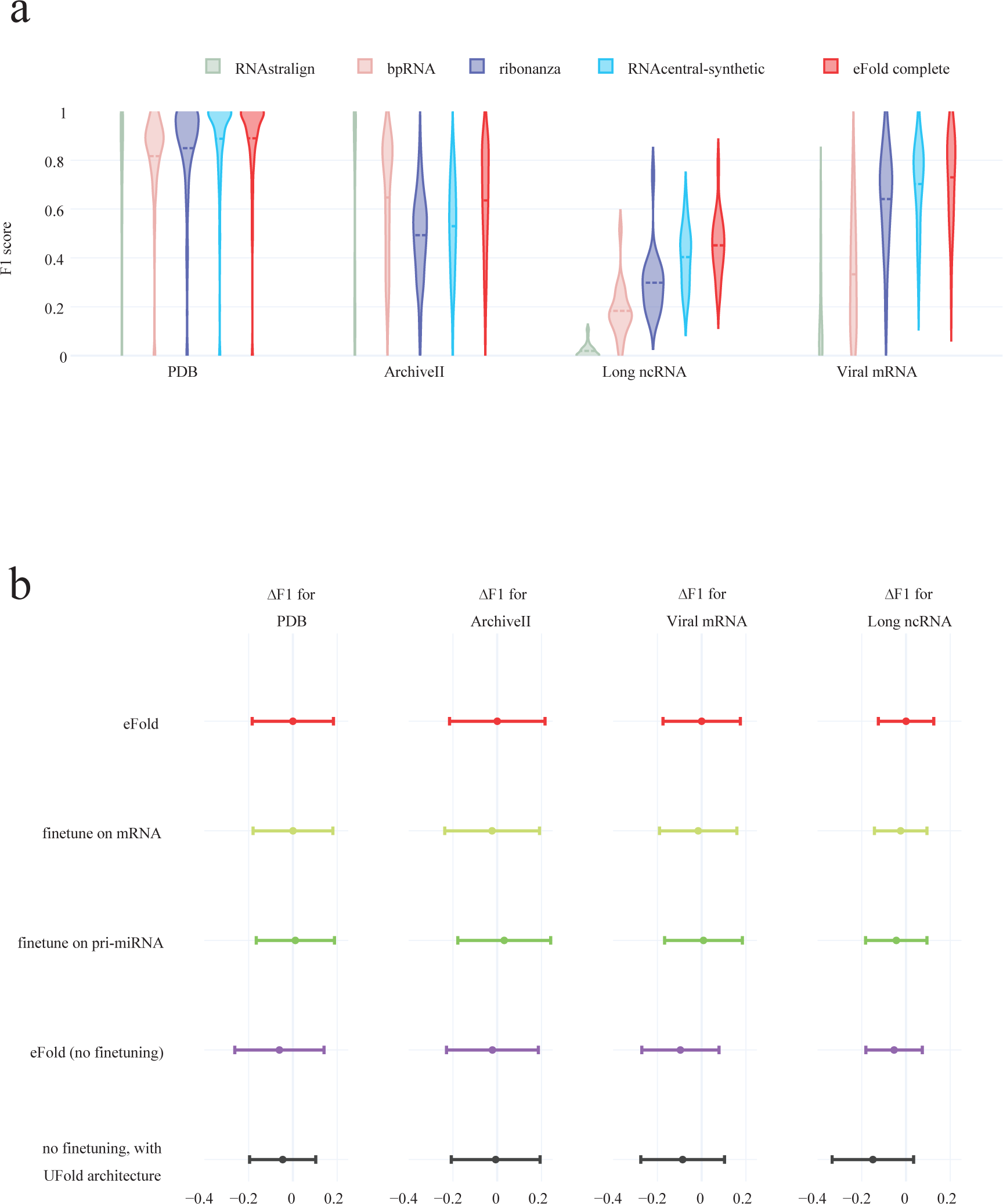
Ablation study. **(a)** F1 distribution on each test set when training the eFold architecture on each database individually. The final eFold model, trained on bpRNA (pink), Ribonanza (blue), and synthetic dataset (light blue) is plotted for reference in red. **(b)** Ablation study for different architecture and training strategies. The baseline model in red is eFold (pre trained then fine-tuned). We try fine-tuning on only the mRNA (light green), only the pri-miRNA (dark green), only pretraining (no fine-tuning, purple), and only pretraining with the UFold architecture. The delta F1 score is with respect to the baseline mean F1 score per dataset. The error bar corresponds to one standard deviation.

## Notes

https://rnandria.org

### Competing Interest Statement

The authors have declared no competing interest.

### Summary of Updates

The last name of the corresponding author was misspelled. We corrected it.

